# Efficient heterologous mRNA production in *E. coli* via protein-facilitated protection

**DOI:** 10.64898/2026.06.15.732261

**Authors:** Zongsheng Zou, Tayyaba Younas, Geoff Dumsday, Victoria S. Haritos, Lizhong He

**Affiliations:** Department of Chemical and Biological Engineering, Monash University, Clayton, VIC, 3800, Australia; CSIRO Manufacturing, Ian Wark Laboratory, Clayton, VIC, 3168, Australia

## Abstract

Messenger RNA (mRNA)-based therapeutics have emerged as a new class of biological medicines, clearly exemplified by the global deployment of mRNA vaccines against the COVID-19 pandemic. Currently, therapeutic mRNA is primarily produced through *in vitro* transcription that suffers high production costs. Until now, intracellular manufacture of mRNA has been challenging due to the presence of ubiquitous RNases *in vivo*. Here, we have developed a new approach that protects eukaryotic mRNA from RNase degradation ensuring longevity and integrity of mRNA inside microbial cells. Through targeted strain and molecular engineering, our approach involves specially designed inserts in mRNA that facilitate formation of stabilized and protected protein-mRNA complexes. In addition to vastly improved stability, the protein-mRNA complexes enable convenient purification of mRNA from cell lysate with high purity using conventional chromatography. The work reported here promises a scalable, rapid, and low-cost approach to produce fully functional eukaryotic mRNA using well-known microbial systems.

## Introduction

As a result of recent technological advances in improving the stability and delivery of mRNA molecules while reducing unfavourable immunogenicity, mRNA-based therapeutics are a rapidly emerging new class of drugs suitable for a range of therapeutic indications(1–4). The COVID-19 pandemic has accelerated the rapid development of mRNA-based technology, with the first COVID-19 mRNA being approved by FDA within a time frame of 10 months from the design of vaccine candidates, a remarkable achievement in history of biological drugs(5,6). Beyond COVID-19 vaccines, mRNA molecules also offer significant advantages for cancer immunotherapy such as safe administration, high potency and a short development cycle(7). After vaccination, injected mRNA is translated into protein in antigen-presenting cells (APCs), facilitating immune stimulation and APC activation(8). With encouraging outcomes from mRNA cancer vaccines established in several clinical trials(9–11), we foresee the rapid advancement of mRNA-based therapy in the field of cancer immunotherapy in the near future.

mRNA therapeutics has several advantages over the current protein and DNA-based therapies(12). The delivery of in vitro transcribed (IVT) coding mRNA results in production of a properly folded and post-translationally modified recombinant protein, avoiding the problems often encountered with current recombinant antibody-based therapeutics: poor stability and protein aggregation during storage(3). More importantly, mRNA-based antibody expression could last for a few days in contrast to the short half-life of recombinant antibodies(13–15). Unlike DNA vectors, mRNA is only biologically functional in cytoplasm, avoiding the risks of gene insertion into the host genome(12). Another significant advantage of mRNA therapeutics is that the mRNA manufacturing process is essentially the same for all RNAs because the physicochemical characteristics of different coding mRNAs are similar. The technology is therefore suitable for rapid response to a pandemic(16) and promising a new therapeutic for personalized medicine(17). Despite these advantages, the current manufacturing of mRNA does have challenges.

At present, mRNA is mainly produced through in vitro transcription (IVT) using phage polymerases, DNA template and ribonucleotide triphosphates (rNTPs), followed by enzymatic capping reaction to produce the final therapeutic mRNA(18). However, due to the notoriously errant activity of RNA polymerase, abortive short transcripts and double stranded RNA (dsRNA) are produced together with target mRNA transcripts(19,20). Since dsRNA impurities have been proven to activate the innate immune response and reduce protein expression from target mRNA transcripts(21,22), total removal of contaminants is one of the key requirements for therapeutic mRNA(23). Extensive chromatographic purification steps are required to remove abortive transcripts and dsRNA efficiently, incurring a large loss of target mRNA during these processes(21). Even though large scale production of mRNA has been achieved, it still largely relies on the utilization of expensive and limited materials(6,24). Hence, there is still an unmet need for the development of low-cost manufacturing processes for clinical grade mRNA production.

The key role of mRNA in normal cellular functions and gene expression involves transcription of genetic information from genomic DNA to mRNA followed by translation into proteins. Thus, it is no surprise that many cellular factors and mechanisms are applied to control/minimize mRNA levels, making the accumulation of heterologous mRNA in vivo challenging(25). The highly structured mRNA molecules generally consist of over a thousand nucleotides with a 5’ untranslated region (UTR), 3’UTR and poly A tail that regulate the functions of mRNA. Molecules of this size and type are prone to rapid degradation by native RNases and therefore difficult to accumulate to any significant levels in vivo. To date, several methods based on naturally stable RNAs or its derivatives such as rRNA(26), tRNAs(27),transfer messenger RNA(28,29) and RNase P RNA(30) have been successfully developed for the production of aptamers and RNA transcripts with less than 350 nucleotides. Specifically, these RNAs are fused with a stabilizing element, which helps target RNA transcripts escape degradation(28). Due to the RNA size limitation of these approaches, in vivo production of heterologous fully functional eukaryotic mRNA is yet to be reported.

Herewith, we have developed a new approach enabling production of fully functional eukaryotic mRNA in *E. coli*. Rationally designed mRNA is stabilized once bound to a recombinant RNA binding protein and this protein also enables efficient capture and purification of target mRNA transcript from cell lysate directly. With our approach, template plasmid DNA and mRNA transcripts can be produced simultaneously in a single *in vivo* production process, without the need to supply the phage polymerases and rNTPs required by *in vitro* transcription processes. To prove the concept, we produced a eukaryotic mRNA encoding super folder green fluorescent protein (sfGFP, 1,471 nucleotides (nt)) and purified target mRNA using conventional chromatography. Finally, we show that the eukaryotic mRNA produced *in vivo* is fully functional by translation into protein using a eukaryotic protein expression system.

## Materials and Methods

### Strain and Plasmid construction

All the plasmids were cloned and propagated in *E. coli* Turbo cells (New England Bio labs Inc.). For protein expression and protein-RNA co-expression, *E. coli* HT115(DE3) (Caenorhabditis Genetics Centre) was used. Target mRNA sequence was compared to a list of RNases binding sites (Supplementary table S1), and potential RNases binding sites were avoided by using alternative codons. The secondary structures of target recombinant mRNA sequences were modelled using the Mfold webserver(31). The target mRNA sequence with correct folding of aptamers (Supplementary Figure S1) and hammer head ribozyme was selected for DNA template synthesis (GenScript Inc.), and T7 promoter and terminator were added to the template for *in vivo* production. The DNA template was cloned to pBAD plasmid (a gift from Michael Davidson, Addgene plasmid # 54762), between SgrAI and NsiI restriction sites. The DNA sequence of the designed protein construct was synthesized (GenScript Inc.) and cloned into the same pBAD plasmid between NdeI and EcoRI, for co-expression of protein and RNA. The DNA template of target mRNA was subcloned in empty pBAD plasmid between EcoRI and XhoI to get the control plasmid.

### *In vitro* transcription

For *in vitro* transcription, the DNA template was prepared by double digestion of the pBAD control plasmid with EcoRI and XhoI restriction endonucleases (New England BioLabs Inc.). After digestion, the DNA template was purified and used to transcribe target mRNA using the T7 High Yield RNA transcription Kit from Vazyme (Vazyme Biotech Co.,Ltd) according to the supplier’s instructions.

### Protein expression and purification

Plasmid containing target protein gene (pBAD_RBP-mCherry-RBP_MTATs-sfGFP) was transformed and expressed in E. coli HT115(DE3) following the heat shock protocol(32). Cell pellet was resuspended in the lysis buffer (50 mM Tris-HCl, 50 mM NaCl, 1 mM EDTA, pH 7.4) and cells were lysed by the probe sonication (Q125 sonicator, Qsonica). After centrifugation, supernatant was clarified with 0.45 µm membrane filter followed by protein purification using immobilized metal affinity chromatography using HisTrap HP column. The binding and washing steps were carried out in 50 mM Tris-HCl, 50 mM NaCl, 1 mM EDTA, pH 7.4 and for elution, the same buffer containing 500 mM imidazole was used. The bound protein was eluted by applying gradient increase of imidazole concentration from 0 mM to 500 mM and the target protein eluted at 500 mM. All the buffers were RNases-free (treated with DEPC).

### Protein RNA co-expression

Cell pellet was resuspended in the lysis buffer (50 mM Tris-HCl, 50 mM NaCl, 1 mM EDTA, pH 7.4) and cells were lysed by the probe sonication (Q125 sonicator, Qsonica). After centrifugation, supernatant was clarified with 0.45 µm membrane filter followed by protein purification using immobilized metal affinity chromatography using HisTrap HP column. The binding and washing steps were carried out in 50 mM Tris-HCl, 50 mM NaCl, 1 mM EDTA, pH 7.4 and for elution, the same buffer containing 500 mM imidazole was used. The bound protein was eluted by applying gradient increase of imidazole concentration from 0 mM to 500 mM and the target protein eluted at 500 mM. All the buffers were RNases-free (treated with DEPC).

### RNA extraction

Cell pellet was resuspended in RNase-free Buffer L (10 mM Tris-HCl, 1 mM EDTA, pH 7.4)(27). An equal volume of saturated phenol (Sigma) was added to the cell suspension. After centrifugation, the aqueous phase was transferred to another tube and supplemented with 0.1 volume of 3 M sodium acetate (pH 5.2) and 3 volumes of absolute ethanol to precipitate RNA. The precipitated RNA was pelleted by centrifugation and washed twice with 75% RNase-free ethanol to remove residual phenol. The RNA pellet was air dried at room temperature for 10 min, dissolved in RNase free water and stored at −80°C for further analysis.

### Denaturing PAGE for RNA

The 3.5% TBE urea gel was casted with 40% acrylamide/bis-acrylamide (19:1) (Bio-Rad Laboratories, Inc.), 8 M urea, and 10xTBE buffer (Bio-Rad Laboratories, Inc.)(33). The gel was run at room temperature for 5 h at 100 V on Mini-PROTEAN Tetra Cell vertical gel electrophoresis system (Bio-Rad Laboratories, Inc.) and stained with SYBR safe DNA gel stain (Thermo Fisher Scientific). Images were taken by Gel Doc XR+ system.

### Reverse transcription

To remove plasmid DNA contamination, 1 U of RNase-free DNase I (Vazyme Biotech Co.,Ltd) was added to 10 µl of RNA sample. The reaction was carried out at 37°C for 15 min and DNase I was deactivated by incubating at 65°C for 10 min. The DNase I digested RNA sample was then used for cDNA synthesis using M-MLV (H-) Reverse Transcriptase (Vazyme Biotech Co.,Ltd), according to manufacturer’s protocol. Oligo dT primer was used for cDNA synthesis. Synthesized cDNA was amplified through PCR with Phanta Flash Super-Fidelity DNA Polymerase (Vazyme Biotech Co.,Ltd). A control PCR reaction was carried out with target gene primers (sfGFP-Fwd: 5’-ATGGCTAAAGGAGAAGAACT, sfGFP-Rev: 5’-TCATTTGTAGAGCTCATCC. Anti-covid nanobody-Fwd: 5’-ATGCAGGTGCAGCTGCAGG, Anti-covid nanobody-Rev: 5’-GTGGCTGCTCACGGTCACC.) to confirm complete removal of DNA contamination(34). The amplified DNA sample was sent for sequencing for further confirmation.

### Agarose gel electrophoresis

Samples were loaded into 1% agarose gel and run in 1xTBE buffer on a Mini-Sub Cell GT Cell electrophoresis unit (Bio-Rad Laboratories, Inc.). The gel was run at constant voltage of 100 V for 50min and stained with SYBR safe DNA gel stain (Thermo Fisher Scientific). Image was taken by Gel Doc XR+ system (Bio-Rad Laboratories, Inc.).

### SDS PAGE

We added SDS/β-mercaptoethanol loading buffer into protein samples and incubated them at 95°C for 5 min before loading on 12% Bolt Bis-Tris precast gels (Life technologies). Electrophoresis was carried out at 120 V for 60 min. Protein bands were stained using microwave procedure of Simply Blue Safe stain (Life technologies).

### Protein-facilitated mRNA purification

Co-expression plasmid (pBAD_RBP-mCherry-RBP_MTATs-sfGFP) was transformed and expressed in *E. coli* HT115(DE3) cells. Cell pellet was lysed through probe sonication (Q125 sonicator, Qsonica), with lysis buffer (50 mM Tris-HCl, 50 mM NaCl, 1 mM EDTA, pH 7.4) prepared with RNase-free water. The cell lysate was centrifuged at 10,000 rpm for 15 min, and supernatant was filtered with 0.45 µm filter. For protein facilitated purification of RNA, HisTrap HP column (GE Lifesciences) was applied to capture protein-RNA complex. The purification was performed using following buffers: buffer A (50 mM Tris-HCl, 50 mM NaCl, 1 mM EDTA, pH 7.4), buffer B (50 mM Tris-HCl, 500 mM NaCl, 500 mM imidazole,1 mM EDTA, pH 7.4), buffer C (50 mM Tris-HCl, 50 mM NaCl, 1 mM EDTA, 8 M urea, pH 7.4) and buffer D (50 mM Tris-HCl, 1 M NaCl, pH 7.4). Buffer A, B and C were prepared with DEPC treated water whereas buffer D was prepared in water containing 1% DEPC. After sample loading, buffer A was applied for washing followed by buffer C wash. Target mRNA was eluted by applying gradient increase of NaCl concentration (buffer D). After mRNA elution, the recombinant RNA binding protein was eluted by applying gradient increase of imidazole concentration (buffer B).

### Cell free expression

For cell-free protein expression, wheat germ extract system (Promega corporation) was used. For 50 µl reaction, 10∼30 µg purified mRNA sample was added as template whereas all other essential reagents were added according to supplier’s instructions. The translation reaction was performed at 30°C for 2 h followed by second incubation at 42°C for 2 h to ensure successful translation as the target mRNA has a complex secondary structure at the 5’ UTR. The expressed protein was then analysed via western blot.

### Western blotting

*The in vitro* translated protein sample was mixed with Bolt LDS sample buffer (Thermo Fisher Scientific) and heated at 70°C for 10 min. The proteins were separated through 10∼12% denaturing polyacrylamide gel and transferred to nitrocellulose membrane by means of a Mini Blot Module (Thermo Fisher Scientific). After transfer, the nitrocellulose membrane was air dried overnight. The dried membrane was then immersed in PBS buffer containing Tween 20, 3% skim milk and anti-GFP (Roche) antibody (Sigma) for 1.5 h and rinsed. Goat anti-Mouse IgG (H+L) secondary antibody (Thermo Fisher Scientific) was then applied with the same condition(35). The membrane was washed three times in PBS buffer containing Tween 20. After washing, 1-Step™ NBT/BCIP substrate solution (Thermo Fisher Scientific) was added to develop the membrane. Image was taken through cell phone.

### Dynamic Light Scattering analysis

Purified RBP-mCherry-RBP (60 µl, 2 mg/mL) was mixed with 60 µl of *in vitro* transcribed mRNA for the assembly. *In vitro* assembled protein-RNA nanoparticles were analysed with Zetasizer Nano-ZS (Malvern Instruments). The size of particles was measured in solution (100 µl) containing free protein, *in vitro* transcribed mRNA or *in vitro* assembled protein-RNA complex. For all samples, three independent measurements with each comprising 10 cycles were performed and particle size distribution was analysed using volume percentage.

### Atomic force microscope imaging

Mica surface (Ted Pella, Inc.) was directly used as matrix without cationization as the mRNA already contained Mg^2+^ providing sufficient electrostatic attraction to the slightly negatively charged mica surface. The protein-RNA nanoparticle sample was diluted and 50 µl sample was loaded onto the freshly cleaved (Scotch taped) mica surface and incubated for 2 minutes at room temperature. Then the surface was rinsed with RNase-free water to remove excess unbound sample(36). The matrix was air dried at room temperature and examined by standard tapping mode in air on a Dimension Icon© atomic force microscope using TESPA-V2 cantilever (Bruker-Nano Corporation). The AFM cantilever spring constant was 42 Nm^-1^ with the resonant frequency in air of 320 KHz and the sample was scanned at a rate of 1 Hz.

## RESULTS

### Unprotected mRNA cannot be produced *in vivo* in *E. coli*

The turnover of intracellular RNA is a fundamental metabolic process, for which RNA degradation is an essential component. As such, in vivo production of heterologous mRNA is extremely difficult due to rapid degradation but there is great interest in improving RNA stability for biotechnology applications(37). Structured sequences at 5’ and 3’ ends of an mRNA sequence can improve its in vivo stability(38). Therefore, we rationalize that “capping” both ends of the heterologous mRNA with aptamers (TmTATsfGFP, Supplementary Table S2) might make stable in vivo mRNA production feasible. To test this hypothesis, we added the TAT RNA aptamer to both ends of mRNA encoding sfGFP (Supplementary Figure 2A). To provide additional protection to mRNA, an RNase III deficient strain of E. coli HT115(DE3) specifically engineered to produce dsRNA(39), was used as production strain alone with conventional *E. coli* BL21(DE3). No target mRNA was recovered from cell lysates, suggesting terminal aptamers alone do not protect mRNA *in vivo*, even within an RNase III deficient strain (Supplementary Figure 2B).

To further prevent the access of exoribonucleases that degrade mRNA in the 5’ to 3’ direction or in the 3’ to 5’ direction, multiple aptamers were placed at both ends of the model mRNA. A three-way junction (3WJ) was fused with the aptamers to assure all aptamers were well folded and displayed(40). As poly A tail is a destabilizing signal in E. coli and often leads to degradation of mRNA(41,42), additional aptamers fused with the 3WJ were placed between the 3’UTR and poly(A) tail (MTATs-sfGFP, Supplementary Table S2) to enhance its protection (Figure 1A). Still, no target mRNA was recovered from *E. coli* BL21(DE3) or HT115(DE3) cell lysates (Figure 1B) indicating that additional protection is needed to produce heterologous mRNA *in vivo*.

**Figure 1:**
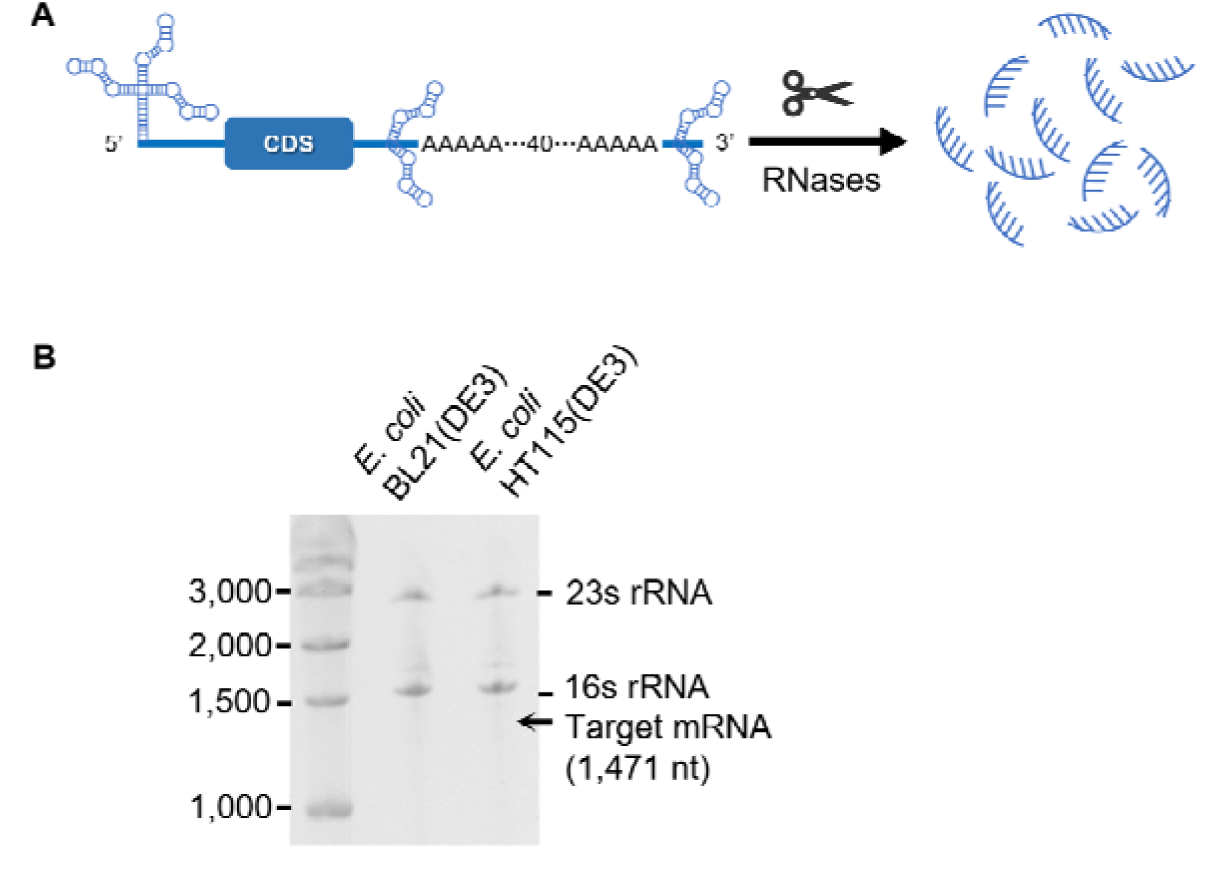
Aptamers alone do not protect mRNA *in vivo*. **A,** Schematic of the mRNA design: multiple aptamers are placed at both ends and 3’ untranslated region of target mRNA. **B,** Target mRNA (MTATs-sfGFP, 1,471 nt) was not observed when expressed in *E. coli* HT115(DE3) and *E. coli* BL21(DE3).

### Recombinant RNA binding protein protect mRNA *in vitro*

Since adding multiple aptamers in combination with an RNase III deficient strain (*E. coli* HT115(DE3)) did not protect the heterologous mRNA from degradation, we explored an additional protection mechanism. As ribosome loading increases mRNA half-life(43), we hypothesized that complexing RNA with proteins may bring steric hindrance to limit RNase access to the heterologous mRNA (Figure 2A). We therefore selected TAT aptamer-binding peptide (RBP, SYGRKKRRQRRRPPQ) to engineer RNA binding protein due to its strong binding affinity with the TAT aptamer (dissociation constant K_d_ = 10^-10^ M)(44). RBPs were fused to both the N- and C-terminus of mCherry (Supplementary Table S3) to generate an RNA binding protein, RBP-mCherry-RBP (Figure 2B). We considered that the interaction between TAT aptamers and the recombinant RNA binding protein was feasible as RBPs are readily accessible to the aptamers (Figure 2C).

**Figure 2:**
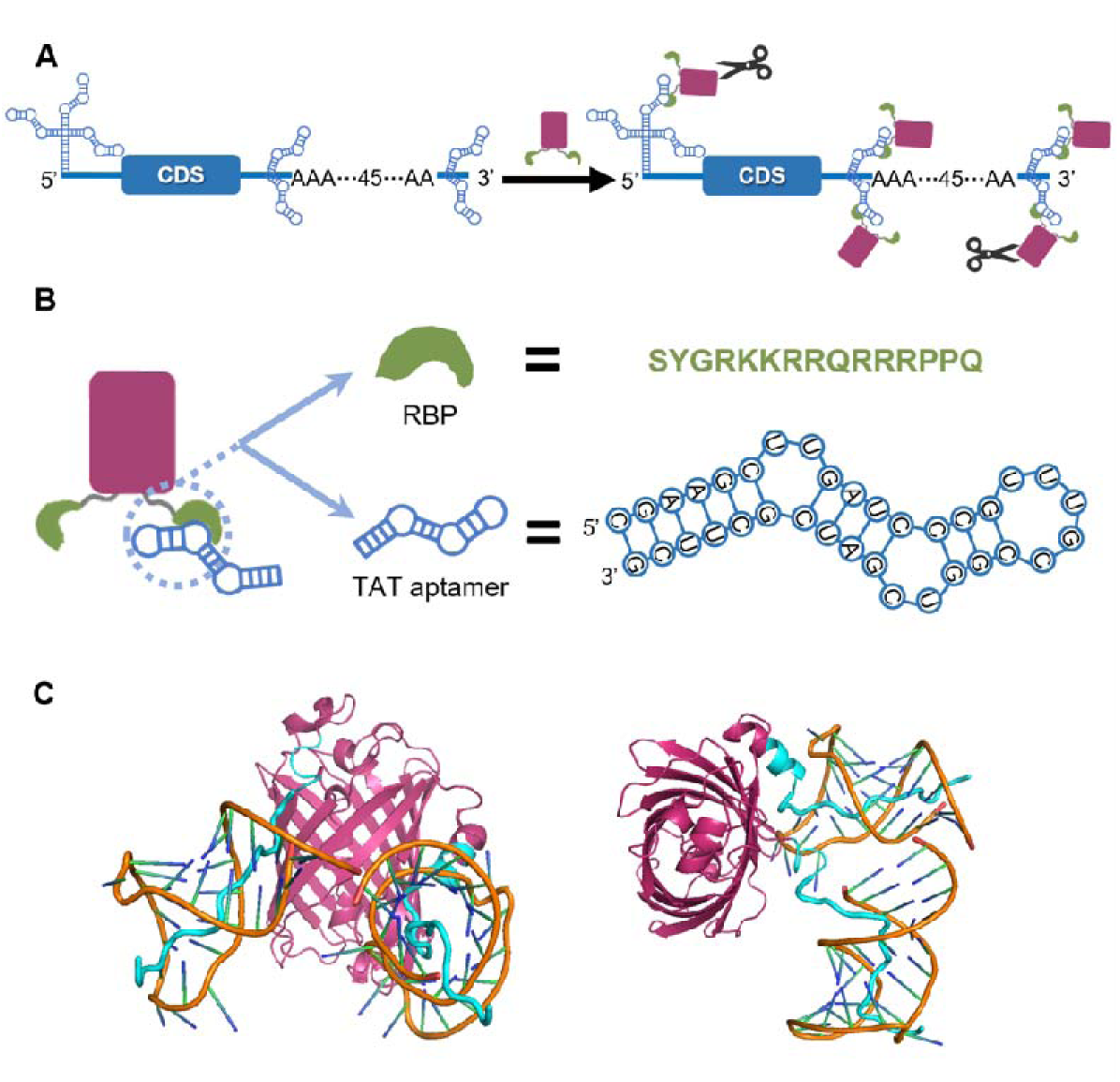
Schematic representation of the designed mRNA sequence and stabilization using a recombinant RNA-binding protein. **A,** Rationally designed mRNA sequence forms particles with recombinant RNA binding proteins and the complex is protected from RNases attack. **B,** Molecular design of mCherry protein fused with aptamer binding peptide with high affinity for TAT aptamer. RBPs are fused to both terminals of mCherry protein. TAT aptamer sequence is also shown. **C,** Schematic showing protein-aptamer interaction (mCherry, pink (PDB: 2H5Q); RBP, cyan; TAT aptamer, orange (PDB: 6MCE)).

The multivalent nature of the binding between mRNA and RBP-mCherry-RBP was expected to result in formation of protein-RNA complex (Figure 3A)). Indeed, dynamic light scattering (DLS) showed that mixing *in vitro* transcribed mRNA with purified RBP-mCherry-RBP protein resulted in the formation of large particles (Figure 3B). Analysed separately, the sizes of *in vitro* transcribed mRNA or purified RBP-mCherry-RBP were less than 50 nm but when mixed, large particles were then detected (two populations, *ca.* 70 nm, and *ca*. 500 nm). The large particles of size around 500 ∼ 700 nm could also be visualized using AFM (Figure 3C). Next, mRNA/protein particles were mixed with clarified lysate from either *E. coli* BL21(DE3) or *E. coli* HT115(DE3) to test resistance of the complex to intracellular RNases. When treated with clarified lysate from *E. coli* BL21(DE3), the complexed RNA was rapidly degraded. In contrast, the mRNA in the complex was stable for at least 5 days when treated with lysate from the RNase III deficient strain, *E. coli* HT115(DE3) (Figure 3D), suggesting our newly designed mRNA-protein complex should accumulate *in vivo* when produced in the RNase III deficient strain.

**Figure. 3:**
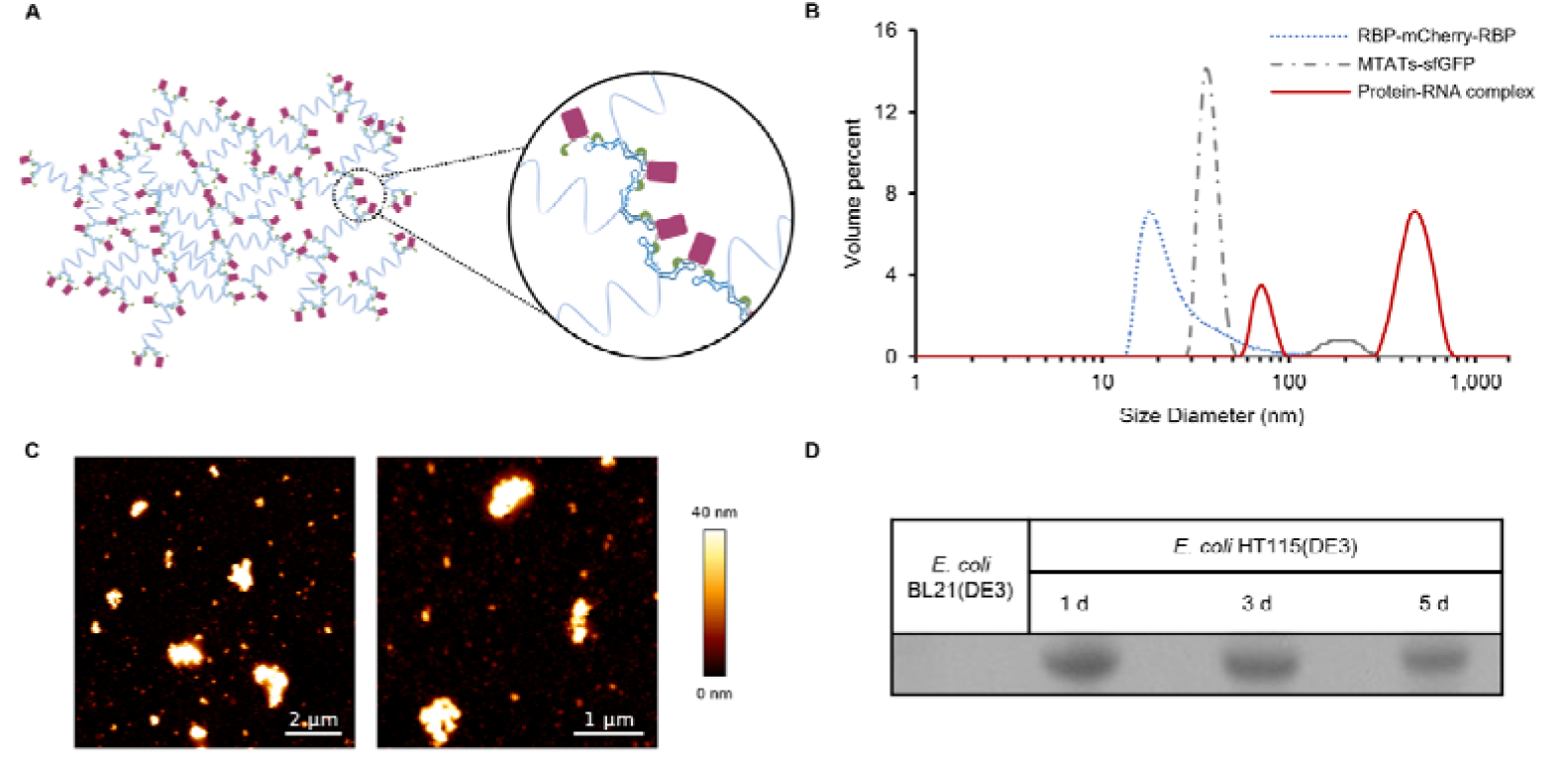
*In vitro* testing of the interaction between RBP-mCherry-RBP and target mRNA. **A**, Schematic representation of the formation of large particles with the engineered mRNA (MTATs-sfGFP, 1,471 nt) and recombinant RNA binding protein. **B**, DLS data showing formation of large particles after mixing RBP-mCherry-RBP with target mRNA. **C**, AFM results confirming the presence of large particles after *in vitro* assembly of RBP-mCherry-RBP and target mRNA sample. **D**, Degradation resistance test showing the protein-RNA complex protects target mRNA from degradation by the RNases in cell lysate from *E. coli* HT115(DE3).

### Production of heterologous mRNA *in vivo*

Co-expression of engineered mRNA and RBP-mCherry-RBP was tested using *E. coli* HT115(DE3) as the host. The pBAD plasmid was modified to enable co-expression of the target mRNA with the gene encoding RBP-mCherry-RBP downstream and regulated by araBAD promoter (Supplementary Figure S3). For efficient transcription, the T7 promoter was used and the designed mRNA sequence started with triple guanine(45,46). Three hours after IPTG induction of target mRNA production, the cells were harvested by centrifugation. Interestingly, the pellet showed a distinct pink colour indicating co-expression of RBP-mCherry-RBP (Figure 4A) even in the absence of arabinose induction, probably due to the low termination efficiency (∼ 80%) of the T7 terminator(47). An mRNA band of 1,471 nt was visualized on 3.5% TBE urea gel (Figure 4B) clearly demonstrating successful production of target sfGFP mRNA. And an mRNA band of 1,147 nt was visualized on 3.5% TBE urea gel (Figure 4D) clearly demonstrating successful production of target anti-covid nanobody mRNA. To confirm the presence of target heterologous mRNA, total RNA was extracted and reverse transcribed into cDNA using an oligo dT primer. The cDNA was amplified using PCR and sfGFP/anti-covid nanobody gene specific primers, with a resultant product of the correct size (Figure 4C and Figure 4E). Sequencing of the PCR product further confirmed the successful production of heterologous mRNA *in vivo* using *E. coli* HT115(DE3).

**Figure 4:**
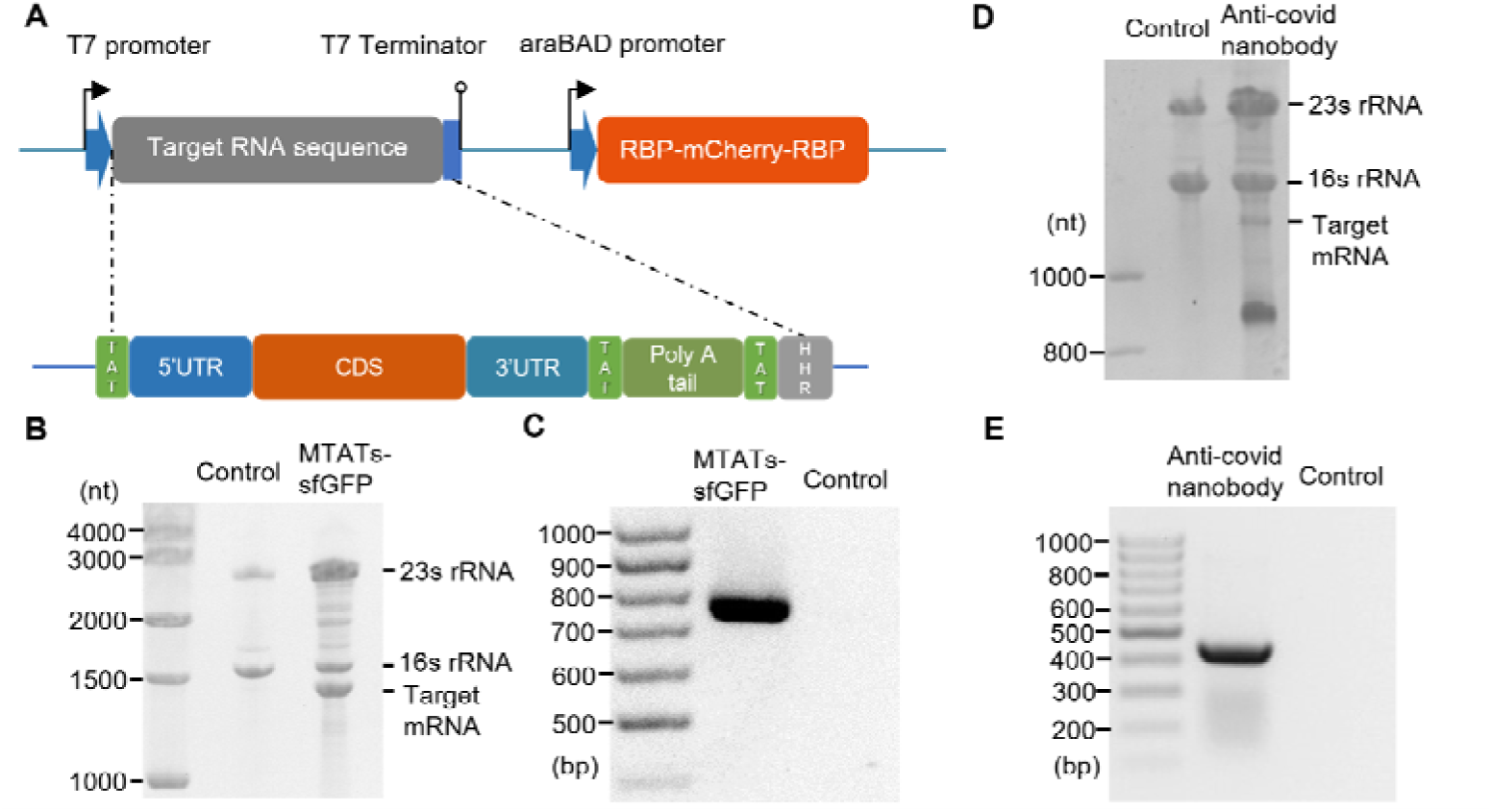
*In vivo* production of heterologous mRNA in *E. coli* HT115(DE3). **A**, Schematic showing the arrangement of genes in the expression plasmid. Target mRNA sequence (MTATs-sfGFP, 1,471 nt) was under the control of the T7 promoter and RBP-mCherry-RBP gene was under araBAD promoter. After IPTG induction, RBP-mCherry-RBP and target mRNA were co-expressed. **B**, 3.5% TBE urea gel results showing production of the MTATs-sfGFP mRNA (1,471 nt). **C**, 1% DNA agarose gel with the product of reverse transcription and PCR from total RNA extracted from *E. coli* HT115(DE3) co-expressing RBP-mCherry-RBP and MTATs-sfGFP mRNA. **D**, 3.5% TBE urea gel results showing production of the MTATs-Anti-covid nanobody mRNA (1,150 nt). **E**, 1% DNA agarose gel with the product of reverse transcription and PCR from total RNA extracted from *E. coli* HT115(DE3) co-expressing RBP-mCherry-RBP and MTATs-Anti-covid nanobody mRNA.

### Protein facilitated purification of target mRNA

Cell lysate containing the protein-RNA complex (with a 6×his tag on the protein) was loaded onto an immobilized metal affinity chromatography (IMAC) resin to capture the protein-RNA complex. Through intensive wash steps, most of the contaminating RNA and proteins from host cells were removed. The target mRNA was eluted from the bound protein-RNA complex using a sodium chloride gradient (buffers contained the RNase inhibitor, diethyl pyrocarbonate). Protein was eluted from the column using imidazole (Figure 5A). Analysis using a 3.5% TBE urea gel (Figure 5B) showed target mRNA was purified through protein facilitated purification with high purity, and RBP-mCherry-RBP was expressed without any apparent truncation (Figure 5C). Note: we included a highly efficient self-cleavage hammerhead ribozyme between the designed mRNA sequence and the T7 terminator (Supplementary Table S2), to assure the homogeneity of produced mRNA sequence(48), which contributed to the high purity of mRNA.

**Figure 5:**
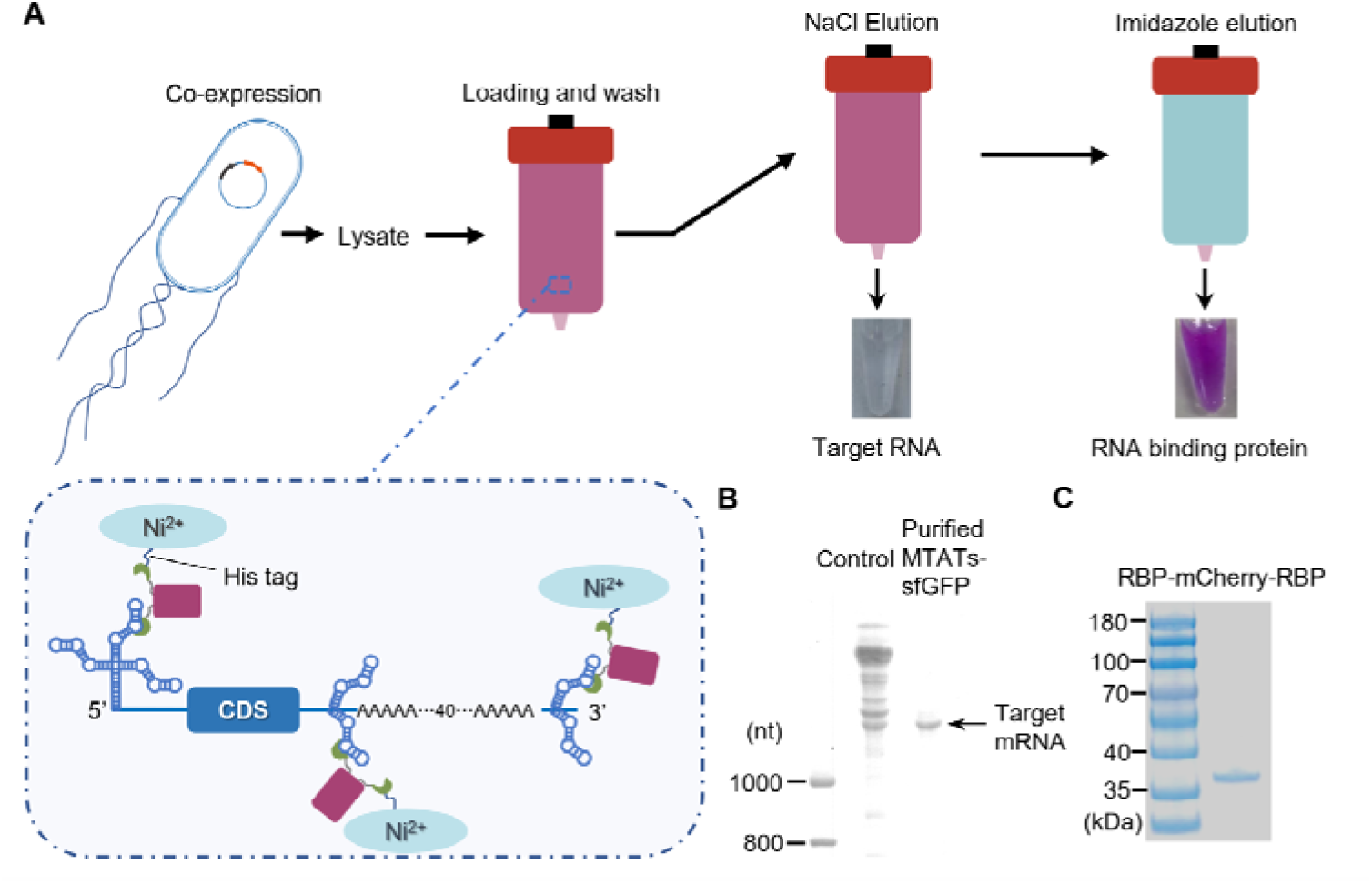
Protein-RNA complex facilitated purification of mRNA produced *in vivo*. **A**, Principle of the purification process. Target mRNA (MTATs-sfGFP, 1,471 nt) complexed with RBP-mCherry-RBP from cell lysate binds to the NTA-Ni resin through a His-tag on mCherry. Target mRNA was eluted with a NaCl gradient elution. **B**, 3.5% TBE urea gel analysis showing target mRNA was successfully purified with high purity. **C**, 12% SDS PAGE result showing that RBP-mCherry-RBP was successfully produced.

### Translation of heterologous mRNA produced *in vivo*

After residual salt was removed from the “IMAC” purified mRNA by ethanol precipitation, a cell-free eukaryotic system, wheat germ extract, was used for *in vitro* translation of the purified mRNA. After *in vitro* translation of target mRNA, fluorescence was detected in the reaction mixture, suggesting production of sfGFP (Figure 6A). The production of sfGFP was further verified via western blot using anti-GFP antibody: a 27 kDa band (the expected size of sfGFP) was detected (Figure 6B) confirming the functionality of the heterologous eukaryotic mRNA produced *in vivo*.

**Figure 6:**
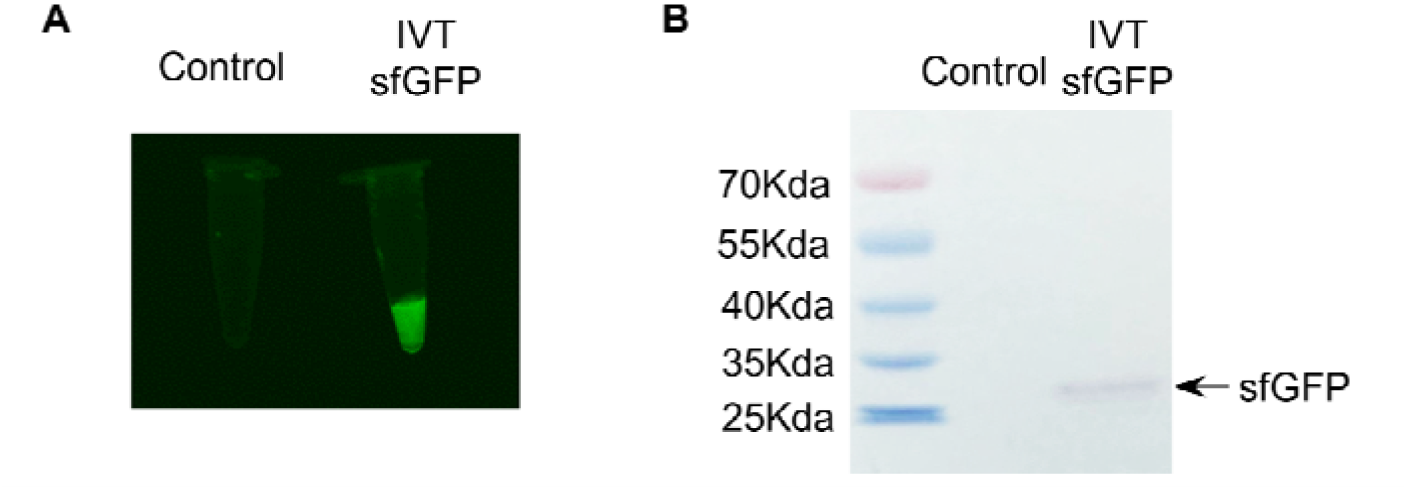
Testing the functionality of heterologous mRNA produced *in vivo*. **A**, mRNA produced *in vivo* was used as a template for *in vitro* translation resulting in production of fluorescent protein. **B**, Western blot confirming production of sfGFP.

## DISCUSSION

In this study, the heterologous mRNA produced in *E. coli* was designed for protein expression using eukaryotic intracellular machinery. The intended application in eukaryotes requires the designed mRNA to have molecular features such as UTR sequences and a poly(A) tail for eukaryotic expression. For example, poly(A) tail plays an essential role in regulation of expression through both stimulating translation and stabilizing mRNA in eukaryotes(49), and the design of mRNA for eukaryotes requires a sufficient length of a poly(A) tail. However, the addition of poly(A) tail often destabilizes mRNA in prokaryotes(50), and RNA degradation in *E. coli* is actually regulated and accelerated by a poly(A) tail(51).The difference in the regulation mechanism between eukaryotes and prokaryotes makes it challenging to produce heterologous eukaryotic mRNA using microbial systems.

As shown in Supplementary Table S4, there are three major mRNA degradation pathways in *E. coli*, and any of them can rapidly degrade heterologous mRNAs. As shown in Figure 1 and Supplementary Figure S2, strategies that tackle some but not all of degradation mechanisms failed to produce detectable target mRNA. We have demonstrated a combined approach that tackles all major degradation mechanisms to efficiently make fully functional eukaryotic mRNA in microbial systems. Specifically, we have added aptamers on the 5’ end which recruit recombinant aptamer binding protein, RBP-mCherry-RBP, to provide sufficient steric hindrance to RNase blocking their access which, in turn, minimizes 5’ to 3’ degradation of mRNA. Similarly, aptamers have been added at the 3’ end to recruit RBP-mCherry-RBP to prevent 3’ to 5’degradation. Particularly, aptamers have been added in front of poly(A) tail to mitigate poly(A) tail dependent 3’-exonucleolytic degradation. In order to mitigate direct entry degradation, we have combined two complementary strategies: 1) careful design of mRNA sequence through selection of different codons to minimize the number of exposed binding sites and thus mitigate cleavage at specific binding sites, and 2) deployment of an RNase III-deficient strain to mitigate cleavage at double-stranded regions. The four strategies summarized in Extended Data Table 1 have led to production of mRNA that has been successfully translated into functional GFP using eukaryotic intracellular machinery.

To the best of our knowledge, it is the first time that *E. coli* has been used to produce fully functional eukaryotic mRNA that contains key features such as UTR sequences and a poly(A) tail for expression using eukaryotic intracellular machinery. Due to the growing needs of large amounts of homogeneous RNA materials for structural, biochemical and pharmacological studies(52), scaffold-based methods have been successfully developed for *in vivo* production of RNA. However, such RNAs are typically less than 350 nt and are not purposely designed for eukaryotic gene expression. In this study, we designed and demonstrated an approach that overcomes three major routes of RNase degradation (Supplementary Table S4) to fully protect heterologous mRNA from degradation in *E. coli*. While the strategies we developed are comprehensive, the execution of the method is straightforward. It only requires a rationally designed mRNA sequence with parallel expression of a recombinant RNA binding protein, RBP-mCherry-RBP, in an RNase III deficient strain, *E. coli* HT115(DE3).

It should be noted that, in *E. coli*, the stability of homologous mRNA largely depends on ribosome loading as transcription is usually coupled with translation(53). However, there is no ribosome loading onto heterologous eukaryotic mRNA in our case because they are designed to be translated, in eukaryotes, not in *E. coli*. Our design of protein binding on the 5’ end can effectively stop exoribonucleases accessing target eukaryotic mRNA from the early stage of transcription. In fact, the removal of the aptamer on the 5’ end results in mRNA degradation (Supplementary Figure S4), confirming the importance of protein binding for protection of mRNA.

Another challenge associated with *in vivo* mRNA production is efficient separation of the target mRNA product from cellular RNAs and proteins with a resulting high-purity product. In this respect, the high binding affinity and specificity of RBP and TAT aptamers facilitates selective capture and efficient purification of target mRNA from cell lysates with a single purification cycle. *In vivo* production has the added advantage: there is no need to produce template DNA, polymerases and rNTPs separately and mix them together. It can be easily scaled for production of a large amount of mRNA at a low cost using existing fermentation technologies and facilities.

In summary, for the first time we have demonstrated production of full-length eukaryotic mRNA within *E. coli*. It is also the first example of a large heterologous RNA transcript (1,471 nt) being successfully produced *in vivo*. This approach also enabled rapid capture and purification of target mRNA directly from cell lysates using conventional chromatographic techniques. Furthermore, we have demonstrated that the *in vivo* produced mRNA is functional in a eukaryotic expression system. Further optimization is needed in future work to reduce the influence of the aptamer sequence in the 5’UTR on translation efficiency, and capping the mRNA may also increase its stability and translation efficiency. Nevertheless, the approach demonstrated here can serve as a solid base for production of fully functional mRNA using microbial systems. Importantly, the production of different target mRNA sequences using our expression system only requires replacement of the coding sequence (CDS), promising a flexible platform to produce mRNA therapeutics in a timely manner at low-cost.

## Data availability

The authors confirm that the data supporting the findings of this study are available within the article and its supplementary materials.

## Supplementary data

Supplementary Data are available at NAR Online.

## Acknowledgements

Z.Z acknowledges the Monash University Office of Graduate Research and ARC Research Hub for Energy-efficient Separation for his Ph.D. scholarship. We acknowledge the assistance of the Monash Proteomic and Metabolomic Facility for protein identification. The authors thank Dr Gina Pacheco Arredondo for assistance in figure preparation and Mr Kevin K.Y. Hu for proof reading of the manuscript.

## AUTHOR CONTRIBUTIONS

L.H. conceived the research. Z.Z. designed mRNAs and plasmids, performed experiments, analysed data and wrote the manuscript. Z.Z. and T.Y. designed mRNA-protein complex. T.Y. conducted AFM experiment and analysis. Z.Z., T.Y., G.D., V.S.H. and L.H. contributed to data analysis and interpretation. G.D., V.S.H. and L.H. supervised the work. All authors reviewed and edited the manuscript.

