## Supplementary Data for "Efficient heterologous mRNA production in *E. coli* via protein-facilitated protection"

### Supplementary Figure S1

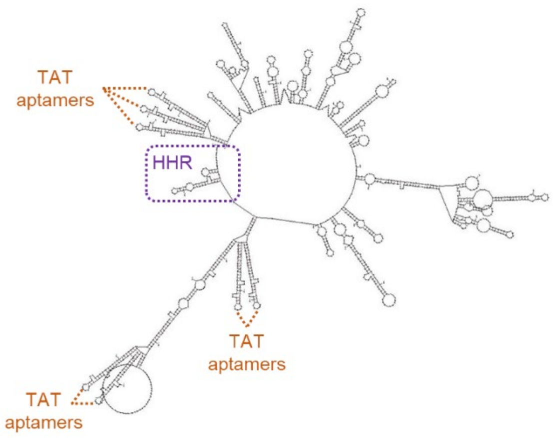


**Supplementary Figure S1. Mfold simulation of the designed eukaryotic mRNA.**

**Supplementary Figure S2**

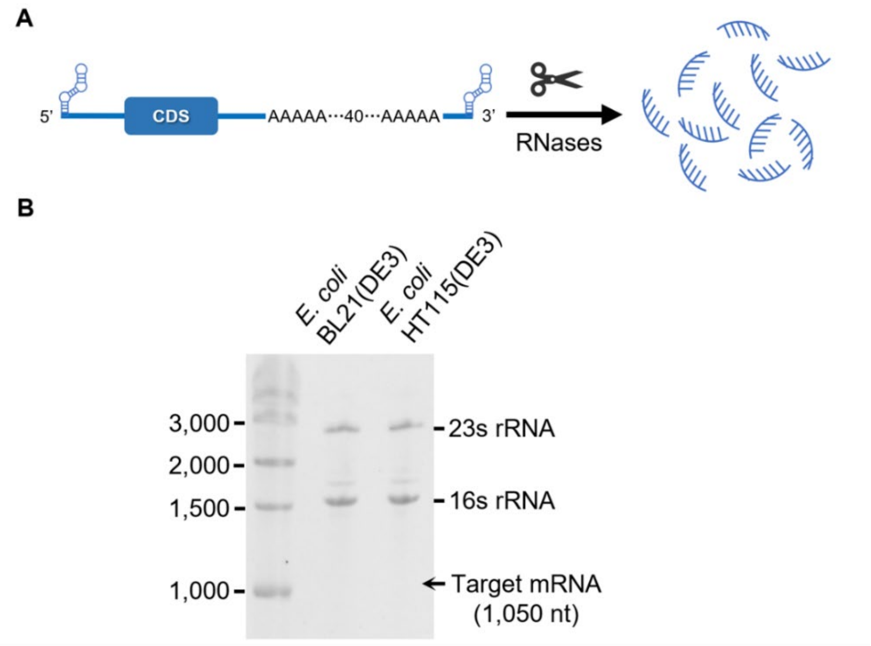


**Supplementary Figure S2. Protection of target mRNA through adding terminal aptamers. A**, Schematic showing the design of target mRNA (TmTATsfGFP) with terminal aptamers. **B**,3.5% TBE gel result showing no target mRNA was detected in cells expressing the target mRNA (1,050 nt).

### Supplementary Figure S3

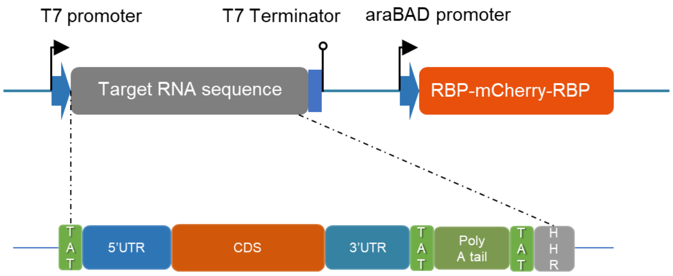


**Supplementary Figure S3. Vector map.** RBP-mCherry-RBP is under the control of araBAD promoter and target RNA is under the control of T7 promoter. Target mRNA contains TAT aptamers and essential components of mRNA: 5'UTR, CDS, 3'UTR, and poly(A) tail. TAT aptamers are placed at 5'UTR and after 3'UTR and poly(A) tail. A hammer head self-cleavage ribozyme is applied to ensure the homogeneity of produced mRNA.

**Supplementary Figure S4**

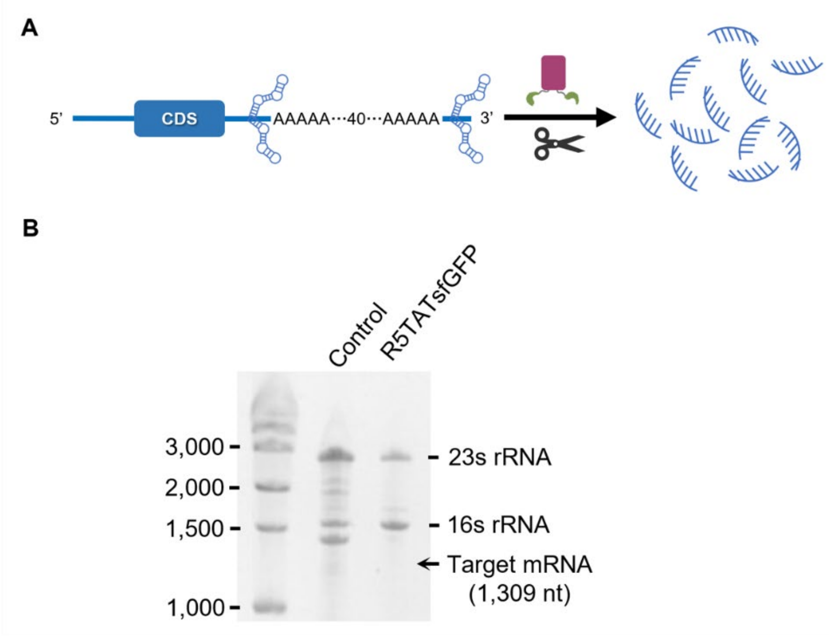


**Supplementary Figure S4. Removal of 5' aptamers leading to mRNA degradation. A**, Schematic showing the design of the target mRNA (R5TATsfGFP) without aptamers on 5' end. **B**, 3.5% TBE gel result showing no target mRNA was detected in cells *E. coli* HT115(DE3) expressing the target mRNA (1,309 nt).

**Supplementary Table S1. Possible cleavage binding sequence of endoribonucleases for direct entry pathway**

| Endoribonucleases | Possible cleavage binding sequence*,** |
|---|---|
| RNase E, RNase G | RNXWUU (with a strong preference for uridine at the +2 position) |
| RNase P | XNNANAU$N_5$U$N_{16}$UU |

* R as G/A, W as A/U, N as any nucleotide and X as the cleavage site.

** See details in the original reference(53)

**Supplementary Table S2. Designed mRNA sequence**

| Name | mRNA size | mRNA sequence* |
|---|---|---|
| MTATs-sfGFP | Before cleavage 1,528 nt<br>After cleavage 1,471 nt | GGGUACGUGAAAGCCGAAGCUUGAUCCCGUUUGCCGGUCGAU<br>CGCUUCGGCUUUCACUCCCGCGCCGAAGCUUGAUCCCGUUUG<br>CCGGUCGAUCGCUUCGGCGCGGAUAUUCGGCCGAAGCUUGA<br>UCCCGUUUGCCGGUCGAUCGCUUCGGCCGAAUAGUACCCCAC<br>AUUUGCUUCUGACACAACUGUGUUCACUAGCAACCUCAAACA<br>GACACCGCCACCAUGGCUGCCACCAUGGCUAAAGGAGAAGAA<br>CUUUUCACUGGAGUUGUCCCAAUUCUUGUUGAAUUAGAUGG<br>UGAUGUUAAUGGGCACAAAUUUUCUGUCCGUGGAGAGGGUG<br>AAGGUGAUGCUACAAACGGAAAACUCACCCUUAAAUUUAUU<br>UGCACUACUGGAAAACUACCUGUUCCAUGGCCAACACUUGUC<br>ACUACUCUGACCUAUGGUGUUCAAUGCUUUUCCCGUUAUCCG<br>GAUCACAUGAAACGGCAUGACUUUUUCAAGAGUGCCAUGCCC<br>GAAGGUUAUGUACAGGAACGCACUAUAUCUUUCAAAGAUGA<br>CGGGACCUACAAGACGCGUGCUGAAGUCAAGUUUGAAGGUG<br>AUACCCUUGUUAAUCGUAUCGAGUUAAAAGGUAUUGAUUUU<br>AAAGAAGAUGGAAACAUUCUCGGACACAAACUGGAGUACAA<br>CUUUAACUCACACAAUGUAUACAUCACGGCAGACAAACAAAA<br>GAAUGGAAUCAAAGCUAACUUCAAAAUUCGCCACAACGUUG<br>AAGAUGGUUCCGUUCAACUAGCAGACCAUUAUCAACAAAAU<br>ACUCCAAUUGGCGAUGGCCCUGUCCUUUUACCAGACAACCAU<br>UACCUGUCGACACAAUCUGUCCUUUCGAAAGAUCCCAACGAA<br>AAGCGUGACCACAUGGUCCUUCUUGAGUUUGUAACUGCUGCU<br>GGGAUUACACAUGGCAUGGAUGAGCUCUACAAAUGAGCUCG<br>CUUUCUUGCUGUCCAAUUUCUAUUAAAGGUUCCUUUGUUCCC<br>UAAGUCCAACUACUAAACUGGGGGAUAUUAUGAAGGGCCUU<br>GAGCAUCUGGAUUCUGCCUAAUAAAAAACAUUUAUUUUCAU<br>UGCGGGUACGUGAAAGCCGAAGCUUGAUCCCGUUUGCCGGUC<br>GAUCGCUUCGGCUUUCACUCCCGCGCCGAAGCUUGAUCCCGU<br>UUGCCGGUCGAUCGCUUCGGCGCGGAUAUUCGGCCGAAGCU<br>UGAUCCCGUUUGCCGGUCGAUCGCUUCGGCCGAAUAGUACCC<br>AAAAAAAAAAAAAAAAAAAAAAAAAAAAAAAAAAAAAAAAA<br>AAAAAAAAAGGGUACGUGAAAGCCGAAGCUUGAUCCCGUUU<br>GCCGGUCGAUCGCUUCGGCUUUCACUCCCGCGCCGAAGCUUG<br>AUCCCGUUUGCCGGUCGAUCGCUUCGGCGCGGAUAUUCGGC<br>CGAAGCUUGAUCCCGUUUGCCGGUCGAUCGCUUCGGCCGAAU<br>AGUACCCGCGCGUCCUGGAUUCGCGGAAACGCGUACAUCCAG<br>CUGACGAGUCCCAAAUAGGACGAAACGCGC |

| | | |
|---|---|---|
| TmTAT sfGFP | Before cleavage 1,107 nt<br>After cleavage 1,050 nt | GGGAAGCUUGAUCCCGUUUGCCGGUCGAUCGCUUCCCCACAU<br>UUGCUUCUGACACAACUGUGUUCACUAGCAACCUCAAACAGA<br>CACCGCCACCAUGGCUGCCACCAUGGCUAAAGGAGAAGAACU<br>UUUCACUGGAGUUGUCCCAAUUCUUGUUGAAUUAGAUGGUG<br>AUGUUAAUGGGCACAAAUUUUCUGUCCGUGGAGAGGGUGAA<br>GGUGAUGCUACAAACGGAAAACUCACCCUUAAAUUUAUUUG<br>CACUACUGGAAAACUACCUGUUCCAUGGCCAACACUUGUCAC<br>UACUCUGACCUAUGGUGUUCAAUGCUUUUCCCGUUAUCCGGA<br>UCACAUGAAACGGCAUGACUUUUUCAAGAGUGCCAUGCCCGA<br>AGGUUAUGUACAGGAACGCACUAUAUCUUUCAAAGAUGACG<br>GGACCUACAAGACGCGUGCUGAAGUCAAGUUUGAAGGUGAU<br>ACCCUUGUUAAUCGUAUCGAGUUAAAAGGUAUUGAUUUUAA<br>AGAAGAUGGAAACAUUCUCGGACACAAACUGGAGUACAACU<br>UUAACUCACACAAUGUAUACAUCACGGCAGACAAACAAAAGA<br>AUGGAAUCAAAGCUAACUUCAAAAUUCGCCACAACGUUGAA<br>GAUGGUUCCGUUCAACUAGCAGACCAUUAUCAACAAAAUACU<br>CCAAUUGGCGAUGGCCCUGUCCUUUUACCAGACAACCAUUAC<br>CUGUCGACACAAUCUGUCCUUUCGAAAGAUCCCAACGAAAAG<br>CGUGACCACAUGGUCCUUCUUGAGUUUGUAACUGCUGCUGGG<br>AUUACACAUGGCAUGGAUGAGCUCUACAAAUGAGCUCGCUU<br>UCUUGCUGUCCAAUUUCUAUUAAAGGUUCCUUUGUUCCCUAA<br>GUCCAACUACUAAACUGGGGGAUAUUAUGAAGGGCCUUGAG<br>CAUCUGGAUUCUGCCUAAUAAAAAACAUUUAUUUUCAUUGC<br>AAAAAAAAAAAAAAAAAAAAAAAAAAAAAAAAAAAAAAAAA<br>AAAAAAAAAGCGAAGCUUGAUCCCGUUUGCCGGUCGAUCGCU<br>UCGCGCGCGUCCUGGAUUCGCGGAAACGCGUACAUCCAGCUG<br>ACGAGUCCCAAAUAGGACGAAACGCGC |

| | | |
|---|---|---|
| MTATs-Anti-covid nanobody | Before cleavage 1,207 nt<br>After cleavage 1,150 nt | GGGUACGUGAAAGCCGAAGCUUGAUCCCGUUUGCCGGUCGAU<br>CGCUUCGGCUUUCACUCCCGCGCCGAAGCUUGAUCCCGUUUG<br>CCGGUCGAUCGCUUCGGCGCGGGAUAUUCGGCCGAAGCUUGA<br>UCCCGUUUGCCGGUCGAUCGCUUCGGCCGAAUAGUACCCCAC<br>AUUUGCUUCUGACACAACUGUGUUCACUAGCAACCUCAAACA<br>GACACCGCCACCAUGGCUGCCACCAUGCAGGUGCAGCUGCAG<br>GAGUCUGGAGGAGGCCUGGUGCAGCCAGGAGGCUCCCUGAGG<br>CUGUCUUGCGCCGUGAGCGGCUUCACCCUGGACUACUAUGCA<br>AUCGGAUGGUUUAGGCAGGCACCAGGCAAGGAGAGGGAGGG<br>CGUGAGCUGUAUCAGCUCCUCUGGCGGCAACACAAAGUACGC<br>AGACUCCGUGAAGGGCCGGUUCACCGCCAGCCGGGACAACGC<br>CAAGAAUACCUUCUACCUGCAGAUGAACAGCCUGAAGCCCGA<br>GGAUACCGCCGUGUACUAUUGCGCCGCCAUCGCCGCCACAUA<br>CUAUUCUGGCAGCUACUAUUUCCAGUGUCCUCACGACGGCAU<br>GGAUUACUGGGGCAAGGGCACCCAGGUGACCGUGAGCAGCCA<br>CGCUCGCUUUCUUGCUGUCCAAUUUCUAUUAAAGGUUCCUUU<br>GUUCCCUAAGUCCAACUACUAAACUGGGGGAUAUUAUGAAG<br>GGCCUUGAGCAUCUGGAUUCUGCCUAAUAAAAAACAUUUAU<br>UUUCAUUGCGGGUACGUGAAAGCCGAAGCUUGAUCCCGUUU<br>GCCGGUCGAUCGCUUCGGCUUUCACUCCCGCGCCGAAGCUUG<br>AUCCCGUUUGCCGGUCGAUCGCUUCGGCGCGGGAUAUUCGGC<br>CGAAGCUUGAUCCCGUUUGCCGGUCGAUCGCUUCGGCCGAAU<br>AGUACCCAAAAAAAAAAAAAAAAAAAAAAAAAAAAAAAAAAA<br>AAAAAAAAAAAAAAAAGGGUACGUGAAAGCCGAAGCUUGAU<br>CCCGUUUGCCGGUCGAUCGCUUCGGCUUUCACUCCCGCGCCG<br>AAGCUUGAUCCCGUUUGCCGGUCGAUCGCUUCGGCGCGGGAU<br>AUUCGGCCGAAGCUUGAUCCCGUUUGCCGGUCGAUCGCUUCG<br>GCCGAAUAGUACCCGCGCGUCCUGGAUUCGCGGAAACGCGUA<br>CAUCCAGCUGACGAGUCCCAAAUAGGACGAAACGCGC |
| R5TAT sfGFP | Before cleavage 1,366 nt<br>After cleavage 1,309 nt | GGGCACAUUUGCUUCUGACACAACUGUGUUCACUAGCAACCU<br>CAAACAGACACCGCCACCAUGGCUGCCACCAUGGCUAAAGGA<br>GAAGAACUUUUCACUGGAGUUGUCCCAAUUCUUGUUGAAUU<br>AGAUGGUGAUGUUAAUGGGCACAAAUUUUCUGUCCGUGGAG<br>AGGGUGAAGGUGAUGCUACAAACGGAAAACUCACCCUUAAA<br>UUUAUUUGCACUACUGGAAAACUACCUGUUCCAUGGCCAACA<br>CUUGUCACUACUCUGACCUAUGGUGUUCAAUGCUUUUCCCGU<br>UAUCCGGAUCACAUGAAACGGCAUGACUUUUUCAAGAGUGCC<br>AUGCCCGAAGGUUAUGUACAGGAACGCACUAUAUCUUUCAA<br>AGAUGACGGGACCUACAAGACGCGUGCUGAAGUCAAGUUUG<br>AAGGUGAUACCCUUGUUAAUCGUAUCGAGUUAAAGGUAUU<br>GAUUUUAAAGAAGAUGGAAACAUUCUCGGACACAAACUGGA<br>GUACAACUUUAACUCACACAAUGUAUACAUCACGGCAGACAA |

| | |
|---|---|
| | ACAAAAGAAUGGAAUCAAAGCUAACUUCAAAAUUCGCCACA<br>ACGUUGAAGAUGGUUCCGUUCAACUAGCAGACCAUUAUCAAC<br>AAAAUACUCCAAUUGGCGAUGGCCCUGUCCUUUUACCAGACA<br>ACCAUUACCUGUCGACACAAUCUGUCCUUUCGAAAGAUCCCA<br>ACGAAAAGCGUGACCACAUGGUCCUUCUUGAGUUUGUAACU<br>GCUGCUGGGAUUACACAUGGCAUGGAUGAGCUCUACAAAUG<br>AGCUCGCUUUCUUGCUGUCCAAUUUCUAUUAAAGGUUCCUUU<br>GUUCCCUAAGUCCAACUACUAAACUGGGGGAUAUUAUGAAG<br>GGCCUUGAGCAUCUGGAUUCUGCCUAAUAAAAAACAUUUAU<br>UUUCAUUGCGGGUACGUGAAAGCCGAAGCUUGAUCCCGUUU<br>GCCGGUCGAUCGCUUCGGCUUUCACUCCCGCGCCGAAGCUUG<br>AUCCCGUUUGCCGGUCGAUCGCUUCGGCGCGGGAUAUUCGGC<br>CGAAGCUUGAUCCCGUUUGCCGGUCGAUCGCUUCGGCCGAAU<br>AGUACCCAAAAAAAAAAAAAAAAAAAAAAAAAAAAAAAAAA<br>AAAAAAAAAAAAAAAAGGGUACGUGAAAGCCGAAGCUUGAU<br>CCCGUUUGCCGGUCGAUCGCUUCGGCUUUCACUCCCGCGCCG<br>AAGCUUGAUCCCGUUUGCCGGUCGAUCGCUUCGGCGCGGGAU<br>AUUCGGCCGAAGCUUGAUCCCGUUUGCCGGUCGAUCGCUUCG<br>GCCGAAUAGUACCCGCGCGUCCUGGAUUCGCGGAAACGCGUA<br>CAUCCAGCUGACGAGUCCCAAAUAGGACGAAACGCGC |

* Orange: TAT aptamers, Deep blue: 5’UTR, Black: target mRNA, Light blue: 3’UTR, Green: Poly A tail, Purple: Hammerhead ribozyme.

**Extended Data Table 3. Amino sequence of RBP-mCherry-RBP.**

| Protein name | Amino sequence* |
|---|---|
| RBP-mCherry-RBP | MHHHHHHGSSYGRKKRRQRRRPPQGSEAAAKMVSKGEEDNMAI<br>IKEFMRFKVHMEGSVNGHEFEIEGEGEGRPYEGTQTAKLKVTKGG<br>PLPFAWDILSPQFMYGSKAYVKHPADIPDYLKLSFPEGFKWERVM<br>NFEDGGVVTVTQDSSLQDGEFIYKVKLRGTNFPSDGPVMQKKTM<br>GWEASSERMYPEDGALKGEIKQRLKLKDGGHYDAEVKTTYKAK<br>KPVQLPGAYNVNIKLDITSHNEDYTIVEQYERAEGRHSTGGMDEL<br>YKEAAAKGSSYGRKKRRQRRRPPQ |

* Pink: Polyhistidine tag, Purple: RNA binding peptide, Light blue: Linker, Red: mCherry

**Supplementary Table S4. mRNA degradation pathways in *E. coli* and design strategies to mitigate intracellular degradation**

| mRNA degradation *in vivo(*38,53) | | | Design strategies to mitigate degradation |
|---|---|---|---|
| Major mRNA degradation pathways | Main RNases involved | Degradation Mechanisms | |
| 5' to 3' degradation | RNase E, RNase J | 5' exonucleases bind to monophosphorylated 5' end and initiate hydrolytic removal of 5'-terminal nucleotides | Addition of aptamers on 5' end and recruiting aptamer binding protein |
| 3' to 5' degradation | RNase II, RNase R, PNPase | poly(A) dependent 3'-exonucleolytic degradation or 3'-exonucleolytic initiation of decay by 3' exonucleases | Addition of aptamers on 3' end and recruiting aptamer binding protein |
| Direct entry | RNase E, RNase G, RNase P | Cleavage at specific binding sites | Sequence designed without exposed binding sites |
| | RNase III | Cleavage at double-stranded regions | RNase III deficient strain |
